# Synaptotagmin-mediated plasma membrane homeostasis is required to recruit the endocytic machinery during cold stress

**DOI:** 10.64898/2026.10.17.759674

**Authors:** José A. Aznar-Moreno, Ángel Del Espino, José Moya-Cuevas, Jessica Pérez-Sancho, Gijs Wilbrink, Michaela Neubergerová, Andreas De Meyer, Steffen Vanneste, Peter Grones, Lourdes Rubio, Roman Pleskot, Daniel Van Damme, Vitor Amorim-Silva, Miguel A. Botella

**Affiliations:** Instituto de Hortofruticultura Subtropical y Mediterránea “La Mayora” (IHSM “La Mayora”), Universidad de Málaga-Consejo Superior de Investigaciones Científicas (UMA-CSIC), Campus Teatinos, 29010 Málaga, Spain; Instituto de la Grasa (IG.CSIC), Edificio 46, Campus Universitario Pablo de Olavide, Ctra. de Utrera Km 1, 41013, Seville, Spain; Cluster of Plant Developmental Biology, Laboratory of Cell and Developmental Biology, Wageningen University & Research, 6708 PB Wageningen, The Netherlands; Institute of Experimental Botany of the Czech Academy of Sciences, Prague, Czech Republic; Department of Experimental Plant Biology, Faculty of Science, Charles University in Prague, Prague, Czech Republic; Department of Plant Biotechnology and Bioinformatics, Ghent University, Technologiepark 71, 9052 Ghent, Belgium; VIB Center for Plant Systems Biology, Technologiepark 71, 9052 Ghent, Belgium; HortiCell, Department Plants and Crops, Faculty of Bioscience Engineering, Ghent University, Ghent, Belgium

## Abstract

Endocytosis depends on plasma membrane (PM) lipid composition, yet how plants sustain endocytic machinery recruitment during stress remains unclear. We investigated whether synaptotagmins SYT1 and SYT3 function in membrane homeostasis and endocytosis in Arabidopsis thaliana. In this work, we revealed that cold stress reversibly blocked FM4-64 internalization in *syt1syt3* double mutants by displacing peripheral and core TPC subunits from the PM. Loss of SYT1 reduced brassinosteroid receptor internalization during cold stress, while double mutants showed enhanced brassinosteroid signaling. *SYTs* and *VAP27* genes showed a condition-dependent genetic interaction in endocytosis: the *syt1/vap27-1/vap27-3* triple mutant resembles *vap27-1/vap27-3* under control conditions and *syt1* during cold stress, suggesting overlapping contributions to endocytic regulation. Using molecular dynamics simulations of protein-membrane interactions, phosphoinositide depletion combined with increased diacylglycerol weakened TPC core membrane association. Our findings identify SYT1 and SYT3 as essential components for sustaining endocytosis during abiotic stress and support a model in which lipid homeostasis at endoplasmic reticulum-PM contact sites preserves the membrane composition required for TPC-dependent endocytosis.

## INTRODUCTION

The plasma membrane (PM) forms a dynamic interface between plant cells and their environment. Its lipid and protein composition must be continuously regulated to maintain cellular integrity while supporting growth, nutrient transport, and responses to external signals (Zhang et al. 2015; Kraus et al. 2024; Zhu et al. 2026). Endocytosis contributes to this regulation by controlling the abundance and distribution of membrane proteins, including receptors and transporters (Arora and Damme 2021; Marhava et al. 2020; Takano et al. 2010). In plants, clathrin-mediated endocytosis is a major route for internalizing PM proteins and is essential for development, hormone signaling, and environmental responses (Arora and Damme 2021; Johnson 2024; Kraus et al. 2024). However, endocytosis itself depends on the properties of the membrane from which vesicles form, creating a need to coordinate membrane homeostasis with endocytic activity (Kraus et al. 2024; Haucke and Kozlov 2018; Posor et al. 2015; Boucrot et al. 2006).

Assembly of the endocytic machinery requires coordinated interactions between cytosolic proteins and membrane lipids. Anionic phospholipids provide binding sites and shape the electrostatic properties of the cytosolic PM surface, with phosphatidylinositol 4-phosphate (PI4P) acting as a major determinant of this electrostatic identity in plants (Simon et al. 2016). Lipid-dependent membrane association is central to the function of the octameric TPLATE complex (TPC), an early and essential adaptor complex in plant clathrin-mediated endocytosis (Kraus et al. 2024). TPC comprises a structured hexameric core and two AtEH/Pan1 subunits, which provide complementary membrane-binding activities. The AtEH/Pan1 subunits interact with phosphatidic acid and phosphatidylinositol 4,5-bisphosphate [PI(4,5)P2] through their Eps15 homology (EH) domains, whereas the hexameric core contains a positively charged membrane-binding interface that favors interactions with PI4P and PI(4,5)P_2_ (Yperman et al. 2021a; Kraus et al. 2025; Yperman et al. 2021b). Membrane-dependent condensation of AtEH/Pan1 is proposed to initiate endocytic assembly by recruiting clathrin and accessory proteins, including adaptor protein complex 2 (AP-2), while the hexameric core promotes membrane bending during vesicle formation (Kraus et al. 2025; Kraus et al. 2024; Dragwidge et al. 2024). These lipid-dependent functions suggest that changes in PM composition could affect both endocytic machinery recruitment and subsequent membrane deformation.

Abiotic stresses challenge this membrane-dependent process. Cold, salinity, and mechanical stress can alter membrane physical properties and activate lipid remodeling and signaling pathways (Guo et al. 2022; Sathasivam et al. 2026; Yu et al. 2021). Although these responses contribute to environmental adaptation, they also create a need to preserve the membrane interactions that support cellular functions. How plant cells accommodate stress-induced lipid remodeling while maintaining the recruitment and activity of endocytic complexes remains poorly understood. Endoplasmic reticulum-PM contact sites (ER-PM CS) provide a link between abiotic stress, lipid homeostasis, and endocytosis at these sites, (Saheki and De Camilli 2017; Benitez-Fuente and Botella 2023). Plant ER-PM CS contain several families of tethering proteins, including synaptotagmins (SYTs), VAMP-associated proteins 27 (VAP27s), and multiple C2 domain and transmembrane region proteins (MCTPs), which contribute to the organization and functional diversity of the ER-PM interface (Morello-López et al. 2026b). Evidence linking these sites to endocytosis comes from VAP27-1 and VAP27-3, which interact with clathrin and are required for normal clathrin dynamics and endocytic trafficking (Stefano et al. 2018).

Among the Arabidopsis SYTs, SYT1 and SYT3 proteins are particularly relevant because of their roles in maintaining PM integrity and lipid homeostasis during abiotic stress (Ruiz-Lopez et al. 2021; Benitez-Fuente and Botella 2023). These ER-anchored proteins contain a synaptotagmin-like mitochondrial lipid-binding protein (SMP) domain and two C-terminal C2 domains, combining lipid-binding capacity with interactions that tether the ER to the PM (Schapire et al. 2008; Benavente et al. 2021). Structural analyses of SYT1 have identified distinct lipid-binding properties of its C2 domains, providing a basis for its regulated association with the PM (Benavente et al. 2021). *SYT1* regulates endosome recycling and movement protein (MP)-mediated trafficking of plant virus genomes (Lewis and Lazarowitz 2010), and SYT1 and SYT3 act redundantly in stress tolerance and prevent excessive diacylglycerol (DAG) accumulation at the PM during cold stress (Ruiz-Lopez et al. 2021). Whether this activity also preserves the membrane environment required for TPC recruitment and endocytosis during stress episodes remains elusive.

Here, we investigate whether SYT1 and SYT3 connect PM lipid homeostasis with endocytosis during abiotic stress. We show that these proteins are required to sustain endocytic activity and TPC association with the PM under cold stress. Our findings support a model in which SYT-mediated lipid homeostasis at ER-PM contact sites maintains a membrane environment permissive for endocytic machinery recruitment.

## RESULTS

### SYT proteins are required for endocytosis under abiotic stress

SYT1 and SYT3 play essential roles in maintaining PM composition and stability under stress (Ruiz-Lopez et al. 2021). Because balanced lipid composition and PM stability are important for clathrin-mediated endocytosis (Boutté and Moreau 2014; Harayama and Riezman 2018), we investigated whether *SYT1* and/or *SYT3* play a role in endocytosis under control and abiotic stress conditions. This was done by quantifying the ratio of intracellular to PM FM4-64 fluorescence as a proxy for overall endocytic flux (Fig. S1), as described (Stefano et al. 2018). Under control conditions, FM4-64 internalization was similar in root cells of Wild-Type (WT) and single *syt1* or *syt3* mutants. In contrast, the *syt1/3* double mutant showed reduced endocytosis compared to WT (Fig. 1a,b). Under cold stress, we observed reduced endocytosis in *syt1*, but not in *syt3*, compared to WT. Remarkably, the double *syt1/3* mutant showed a complete block of FM4-64 internalization during cold stress (Fig. 1c,d), but resumed endocytosis 30 min after transfer to control conditions, confirming that cells remain viable (Fig. 1e,f). Next, we used brefeldin A (BFA) to block endosomal recycling and facilitate BFA body formation to enhance monitoring of endocytic trafficking (Geldner et al. 2001; Stefano et al. 2018). *syt1/3* contained reduced BFA body formation in control conditions compared to WT, and it was absent under cold stress, whereas WT produced BFA bodies under both conditions (Fig. 1g-j). To investigate whether defective syt1/3 endocytosis was specific to cold stress, we analyzed internalization of the FM4-64 *syt1/3* mutant under salt stress. NaCl treatment also reduced endocytosis in *syt1/3* compared to WT, but to a lesser extent than cold, indicating that other stresses also required SYT1 and SYT3 for endocytosis to proceed normally (Fig. S2).

**Figure 1.**
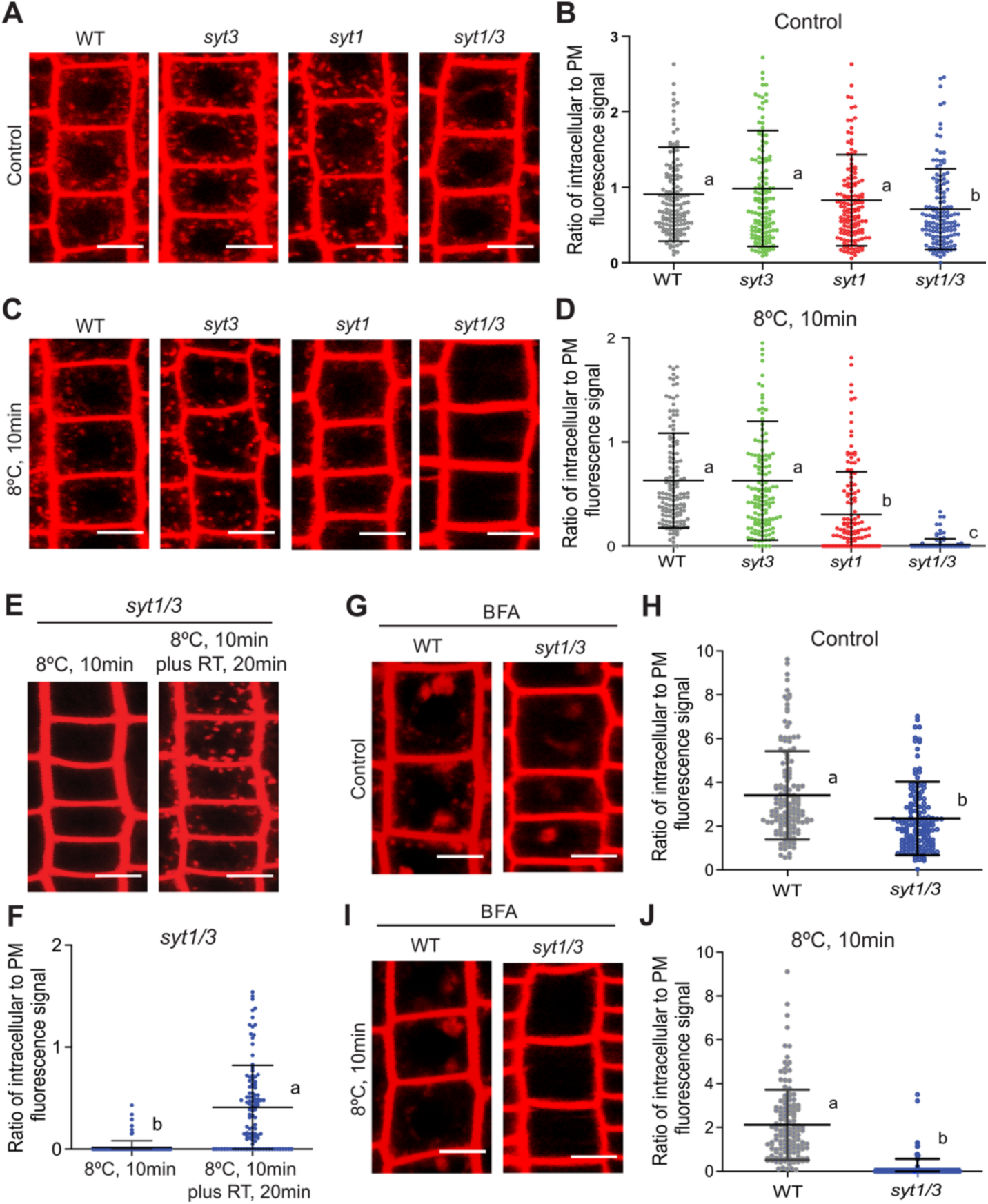
Short-term cold treatment block endocytosis in *syt1/3* double mutants. (**A – D**) Representative confocal images and quantification of FM4-64 internalization into endosomes under control and short-term cold treatment in 6-day old WT, *syt3*, *syt1* and *syt1/3* root epidermal cells. Individual dots represent cells. The lines represent the mean and the standard deviation (SD). One-way ANOVA analysis of variance with Tukey’s Multiple comparison test with *p*<0.05 and n>140 cells. **(E and F)** Representative images and quantification that endocytosis recovers in *syt1/3* mutants following cold treatment. Endocytic flux was determined under cold stress by staining with FM4-64 at 10 min and following a 20 min recovery period at room temperature. Individual dots represent cells. The lines represent the mean and the standard deviation (SD). One-way ANOVA analysis of variance with Tukey’s Multiple comparison test with *p*<0.05 and n>100 cells **(G – J)** Representative confocal images and quantification of FM4-64 internalization into BFA bodies under control and short-term cold treatment in 6-day old WT and *syt3*, *syt1* and *syt1/3* root epidermal cells. Seedlings were exposed to BFA (50 µm) for 50 min following with FM4-64 staining for 10 min. Individual dots represent cells. The lines represent the mean and the standard deviation (SD). One-way ANOVA analysis of variance with Tukey’s Multiple comparison test with *p*<0.05 and n>140 cells. For all assays, at least nine individual seedlings for each line were analyzed. Different letters indicate statistically significant differences. Scale bars, 10 µm.

### SYT1 and SYT3 are required for cargo internalization under stress

Brassinosteroids (BRs) and their PM receptor BR INSENSITIVE1 (BRI1) are among the best-characterized receptor-ligand systems for studying the relationship between receptor endocytosis and plant signaling. The PM-localized pool of BRI1 primarily governs BR signaling, while BRI1 endocytosis attenuates receptor signaling (Irani et al. 2012; Vilarrasa-Blasi et al. 2014). Therefore, we analyzed the effect of *a SYT1* mutation on endocytic cargo by quantifying the distribution between the Trans-Golgi Network (TGN) and the PM of a functional BRI1:BRI1-GFP construct in *syt1* (hereafter *syt1*/BRI1-GFP) and WT backgrounds, using the intracellular-to-PM BRI1-GFP signal ratio (Irani et al. 2012). As with FM4-64, we found no differences in intracellular/PM fluorescence between BRI1-GFP and *syt1*/BRI1-GFP under control conditions (Fig. 2a,b). However, under cold stress, the intracellular/PM BRI1-GFP ratio increased in WT but not in the *syt1* background. This supports the hypothesis that, under cold stress, BRI1-GFP accumulates at the PM in the *syt1* background due to reduced endocytosis (Fig. 2a,b).

**Figure 2.**
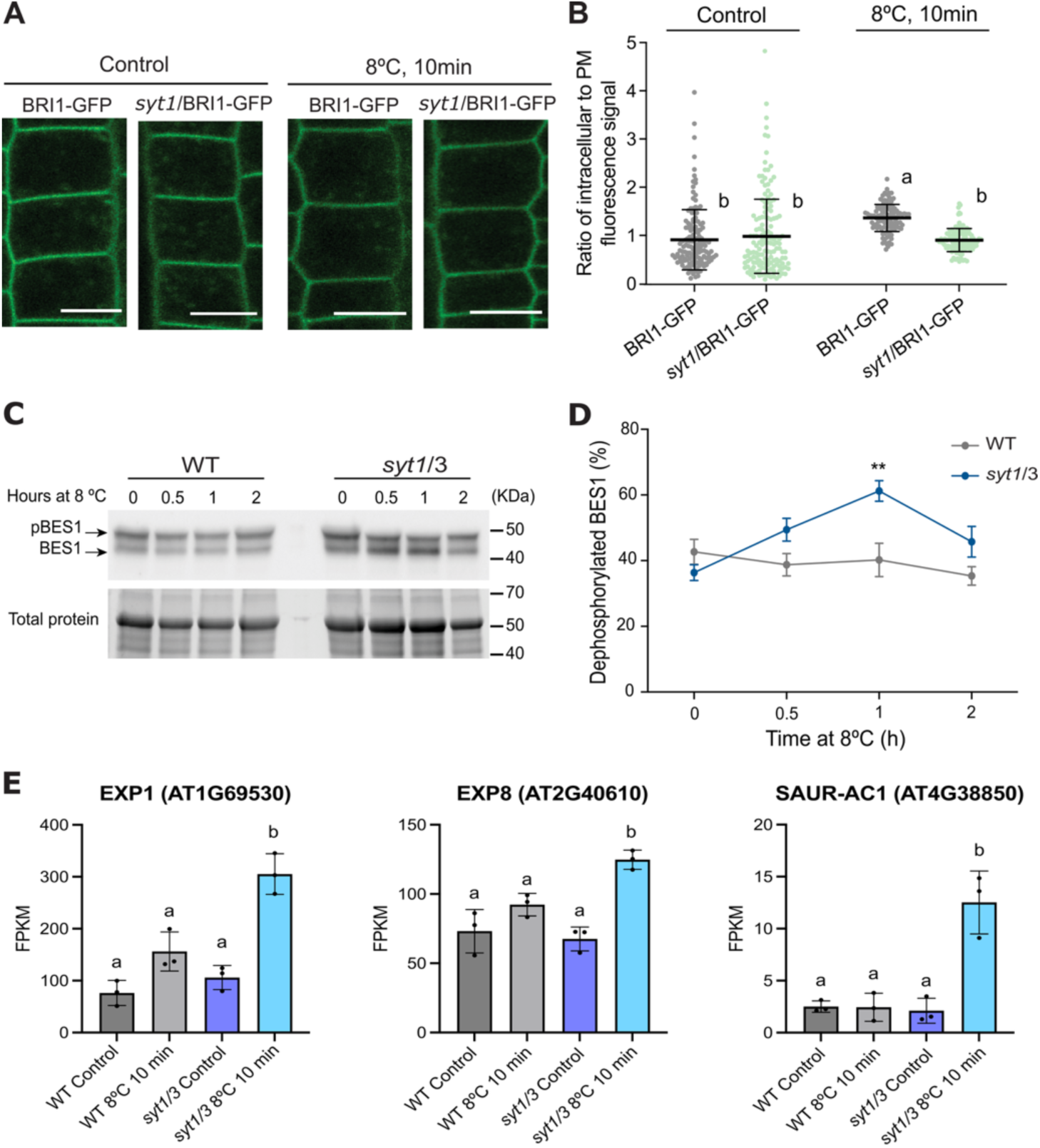
Internalization of BRI1 cargo is delayed in *syt1* single mutant. (**A and B**). Representative confocal images and quantification of the fluorescence intensity of BRI1:BRI1-GFP in PM of 6-days old WT and *syt1* mutants under control and short-term cold treatment. Individual dots represent cells. The lines represent the mean and the standard deviation (SD). Two way-ANOVA with Tukey’s multiple comparison test and *p*<0.05, n>100 cells. For all assays, at least nine individual seedlings for each line were analyzed. Different letters indicate statistically significant differences. Scale bars, 10 µm. **(C and D)** Immunoblot analysis and quantification monitoring the phosphorylation status of BES1 in control conditions and in response to cold treatment for 30 min, 1h and 2h in WT and *syt1/3* mutant. Total proteins were analyzed by an immunoblot assay with a specific anti-BES1 antibody (Yu et al. 2011). The top band corresponds to pBES1 and the bottom band to dephosphorylated BES1 (BES1). N=3 biological replicates, each biological replicate represents a pool of 20 seedlings. Two way-ANOVA with Šidák’s multiple comparison test and *p* <0.05, n=3 independent experiments. Error bars represent SEM. **(E)** Higher expression of BR-induced genes is observed in double mutant *syt1/3* under cold stress. The Fisher’s exact test with the False Discovery Rate (FDR) correction as calculated by the Benjamini-Hochberg procedure was used to calculate statistical differences, FDR<0.05. Error bars represent standard deviation. Different letters indicate statistically significant differences.

Given that PM accumulation of BRI1 activates BR signaling (Irani et al. 2012), we analyzed dephosphorylation of BRI1-ETHYL METHANESULFONATE SUPPRESSOR1 (BES1) as a proxy of increased signaling caused by impaired endocytosis of BRI1 (Belkhadir and Jaillais 2015; Wang et al. 2014). To this end, we analyzed the phosphorylation status of BES1 under control conditions and after 30 min, 1 h, and 2 h of cold stress in WT and *syt1/3* seedlings.

Cold stress caused increased dephosphorylation of BES1 in *syt1/3* compared to WT (Fig. 2c,d), indicating enhanced BR signaling in *syt1/3* compared to WT plants, likely caused by accumulation of BRI1 at the PM.

To investigate molecular responses to *SYT1* and *SYT3* mutations, we identified differentially expressed genes (DEG) in *syt1/syt3* compared to WT seedlings under control conditions and after 10 min of cold treatment at 8 °C, conditions that block endocytosis in *syt1/3* (Fig. S1, S3a,b). Under control conditions, 101 DEG (39 up– and 62 down-regulated) (FDR < 0.05, Log_2_ fold change > 1) were identified in *syt1/3* relative to WT (Fig. S3b,c). Gene ontology (GO) analysis revealed enrichment in jasmonate and wounding responsive genes among the *syt1/3-* induced genes. This indicates that *syt1/3* shows basal stress responses without cold (Fig. S3d), consistent with the slight endocytosis defects observed in the *syt1/3* mutant in control conditions (Fig. 1a,b). In *syt1/3*, cold stress caused large changes in gene expression relative to WT, with 910 differentially expressed genes – DEGs (258 up– and 652 down regulated) (Fig. S3b,c). GO analysis indicated that induced genes were involved in responses to lipid stress, while the repressed genes were mostly housekeeping genes (Fig. S3d). These results support that cold-induced inhibition of *syt1/3* endocytosis causes PM instability, which in turn triggers lipid-derived danger signals, including jasmonic acid biosynthesis, and downstream defense responses.

Using this database, we analyzed the expression of the BR-regulated genes EXPANSIN A1 (EXP1), EXPANSIN A8 (EXP8), and SMALL AUXIN UP RNA 1 (SAUR-AC1) in WT and *syt1/3* seedlings using the RNA-seq data. Cold stress led to higher expression of these genes in *syt1/3* compared to WT plants, consistent with increased BR signaling caused by a PM accumulation of BRI1 (Fig. 2e) (Irani et al. 2012).

### *SYT1* associates with proteins with a role in endocytosis

The ER-PM CS tethers VAP27 proteins, interact with clathrin and are required for clathrin dynamics at endocytic membranes and for endocytic integrity (Stefano et al. 2018). Using a recent study from our group reporting SYT1 associated proteins (Morello-López et al. 2026a), we identified VAP27 and RAB GTPase family proteins involved in endomembrane trafficking (Ebine et al. 2011; Minamino and Ueda 2019) (Fig. 3a). Using an Arabidopsis VAP27-GFP line we confirmed the association between endogenous SYT1 and VAP27-1 by coimmunoprecipitation (Co-IP) (Fig. 3b). As a positive control we used a line expressing SYT5-GFP (García-Hernández et al. 2025) and as a negative control we used an Arabidopsis line expressing free GFP (Fig. 3b). This is consistent with the high colocalization of SYT1 and VAP27-1 at ER-PM CS (Siao et al. 2016; Pérez-Sancho et al. 2015). To investigate if SYT1 and VAP27 interact in vivo, we used Förster Resonance Energy Transfer (FRET). However, SYT1-GFP and VAP27-mCherry do not appear to directly interact at the ER-PM CS, while SYT1-GFP and SYT1-mCherry showed interaction as expected since SYT1 forms homodimers through their SMP domain (García-Hernández et al. 2025; Benitez-Fuente and Botella 2023) (Fig. 3c). This is consistent with the high enrichment VAP27 in the interactome in the APM-MS, but not in the TurboID (Morello-López et al. 2026a), and reinforcing that SYT1 and VAP27 proteins are part of a large complex at ER-PM CS without necessarily being in a sufficiently close proximity for efficient TurboID labeling.

**Figure 3.**
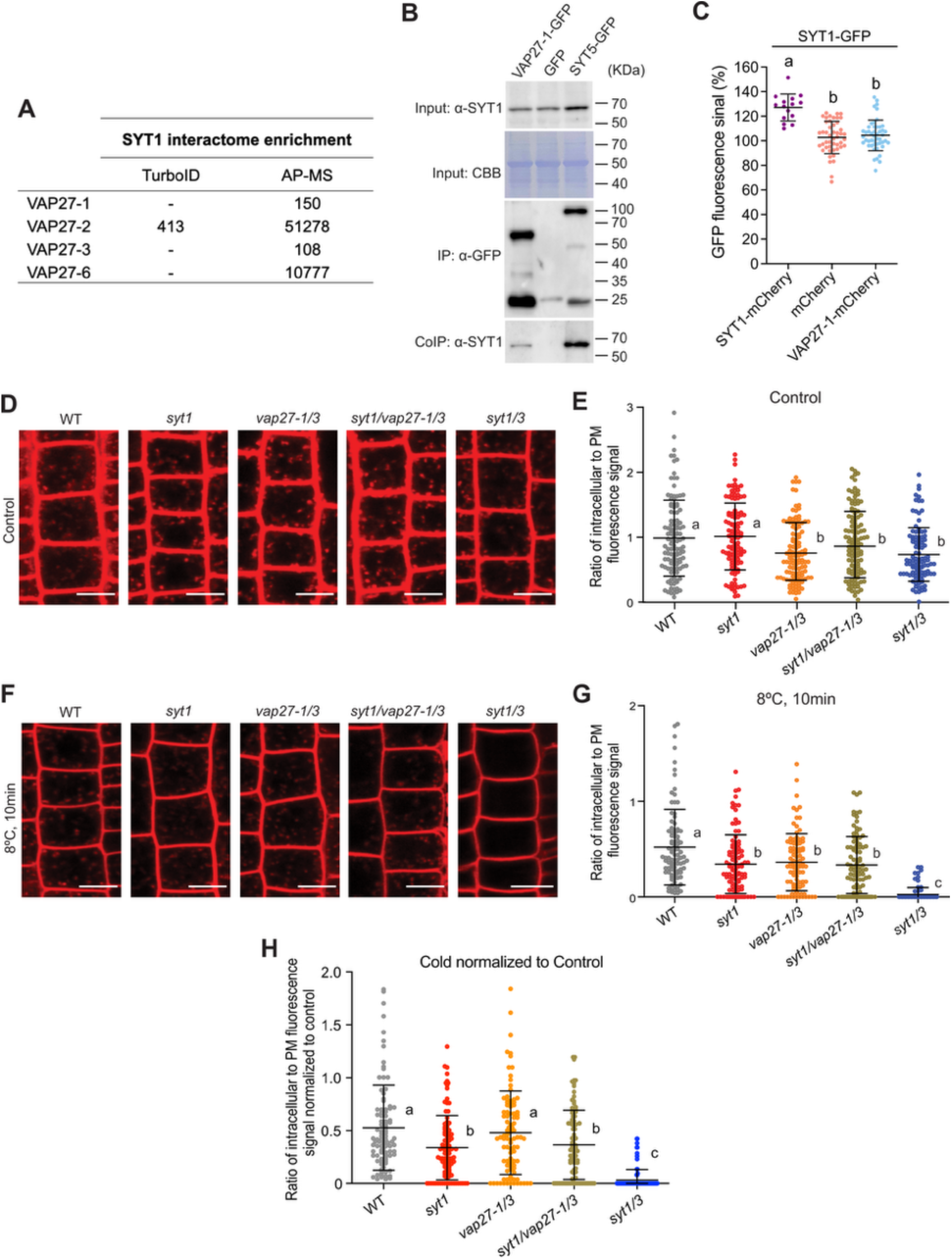
*SYT1* is associated to VAP27 *in vivo*. **(A)** Affinity Enrichment Scores (AES) of VAP27 protein family peptides from Turbo-ID and AP-MS proteomic experiments (Morello-López et al. 2026a). **(B)** Co-immunoprecipitation of SYT1 in 6-day-old seedlings of the stable transgenic VAP27-1-GFP Arabidopsis line. A SYT5-GFP and a free-GFP stable transgenic lines were used a SYT1 positive and negative interactors, respectively. Total (input), IP, and CoIP fractions were analyzed by immunoblotting. Equal loading was confirmed by Coomassie blue staining (CBB) of input samples. SYT1 was detected with an anti-SYT1 antibody. **(C)** FRET analysis showing that SYT1 does not directly interact with VAP27-1. Co-expression of SYT1-GFP with SYT1-mCherry served as a positive control, whereas co-expression with free mCherry was used as a negative control. An increase in GFP fluorescence intensity following mCherry photobleaching was observed exclusively in the positive control (SYT1-GFP / SYT1-mCherry), indicating FRET occurrence. In contrast, no significant increase in GFP signal was detected upon co-expression with VAP27-1-mCherry or free mCherry, demonstrating a lack of direct interaction between SYT1 and VAP27-1. One way-ANOVA with Tukey’s multiple comparison test and *p*<0.05, n>15 assays. **(D – G)** Double *syt1/3* mutant, but not triple *syt1/vap27-1/3* mutant, shows a temporarily block of the endocytosis after short-term cold treatment. Representative confocal images and quantification of FM4-64 internalization into endosomes under control and short-term cold treatment in 6-day old WT, *syt1, vap27-1/3, syt1/vap27-1/3* and *syt1/3* mutants root epidermal cells. Individual dots represent cells. The lines represent the mean and the standard deviation (SD). One-way ANOVA analysis of variance with Tukey’s Multiple comparison test with *p*<0.05 and n>100 cells. **(H)** Normalization of the endocytic flux in control conditions and after short-term cold treatment from panel E and G. Individual dots represent cells. The lines represent the mean and the standard deviation (SD). One way-ANOVA with Tukey’s multiple comparison test and *p*<0.05, n>92 cells. For all assays, at least nine individual seedlings for each line were analyzed. Different letters indicate statistically significant differences. Scale bars, 10 µm.

Our data indicate that SYT1 and SYT3 function in endocytosis, as previously shown for the ER-PM CS tethers VAP27-1 and VAP27-3 (Stefano et al. 2018). To investigate their genetic interactions, we generated a *syt1/vap27-1/vap27-3* triple mutant (hereafter *syt1/vap27-1/3*) by crossing. We did not include the *syt1/syt3/vap27-1/vap27-3* quadruple mutant because the severe endocytic defects of *syt1/syt3* could obscure additional effects of VAP27 loss. Under control conditions, *syt1* showed no detectable endocytic defects, whereas the triple mutant showed reduced endocytosis comparable to *vap27-1/3* (Fig. 3d,e). Under cold stress, the triple mutant instead resembled *syt1* (Fig. 3f,g). Thus, the combined loss of SYT1 and VAP27-1/3 did not exacerbate the defects observed in the affected parental mutant under either condition, consistent with a non-additive, condition-dependent genetic interaction. This pattern suggests that loss of SYT1 masks VAP27-1/3 contribution to the endocytic phenotype during cold stress. Comparison of endocytic activity under cold stress relative to control conditions (Fig. 3h), together with our analysis of SYT mutants, supports a role for VAP27-1/3 in basal endocytosis and an additional requirement for SYT1 and SYT3 to sustain endocytosis during abiotic stress.

### SYT1 and SYT3 are required for TPLATE complex recruitment to the plasma membrane during cold stress

*SYT1* and *SYT* are essential for endocytosis, particularly under stress conditions. Next, we aimed to investigate the localization of TPC components in plant clathrin-mediated endocytosis in a *syt1* and *syt1/3* double mutant relative to WT in control and under cold stress (Kraus et al. 2024). We first investigated the localization of endogenous AtEH1/Pan1 using immunofluorescence localization in Arabidopsis roots using specific antibodies (Grones et al. 2022). As previously described (Grones et al. 2022), confocal imaging revealed a punctate staining pattern at the PM of AtEH1/Pan1 in WT seedlings during control conditions, similar to that of *syt1*, and *syt1/3* seedlings (Fig. S4). After cold stress (8°C, 10 min), WT and *syt1* mutants retained AtEH1/Pan1 signal at the PM, while very little PM signal remained in the *syt1/3* mutant (Fig. S4), indicating that membrane recruitment of the peripheral AtEH1/Pan1 TPC subunits was impaired under cold.

To determine whether the loss of SYT1 and SYT3 also affects PM recruitment of core TPC subunits, we used functional TPLATE-GFP and TML-GFP fusions expressed in their respective mutant backgrounds (Van Damme et al. 2006; Gadeyne et al. 2014). We introduced mutations in *SYT1* and *SYT3* into these reporter lines using CRISPR/Cas9, with two guide RNAs designed for each gene (Fig. 4a and Fig. S5a,b). T2 plants lacking detectable SYT1 protein were identified by immunoblotting and subsequently screened by sequencing for loss-of-function mutations in *SYT3* (Fig. S5a-d). We designated the resulting lines *syt1-3crispr/syt3-4crispr* TPLATE-GFP and *syt1-4crispr/syt3-5crispr* TML-GFP, hereafter *syt1/3crispr* TPLATE-GFP and *syt1/3crispr* TML-GFP, respectively.

**Figure 4.**
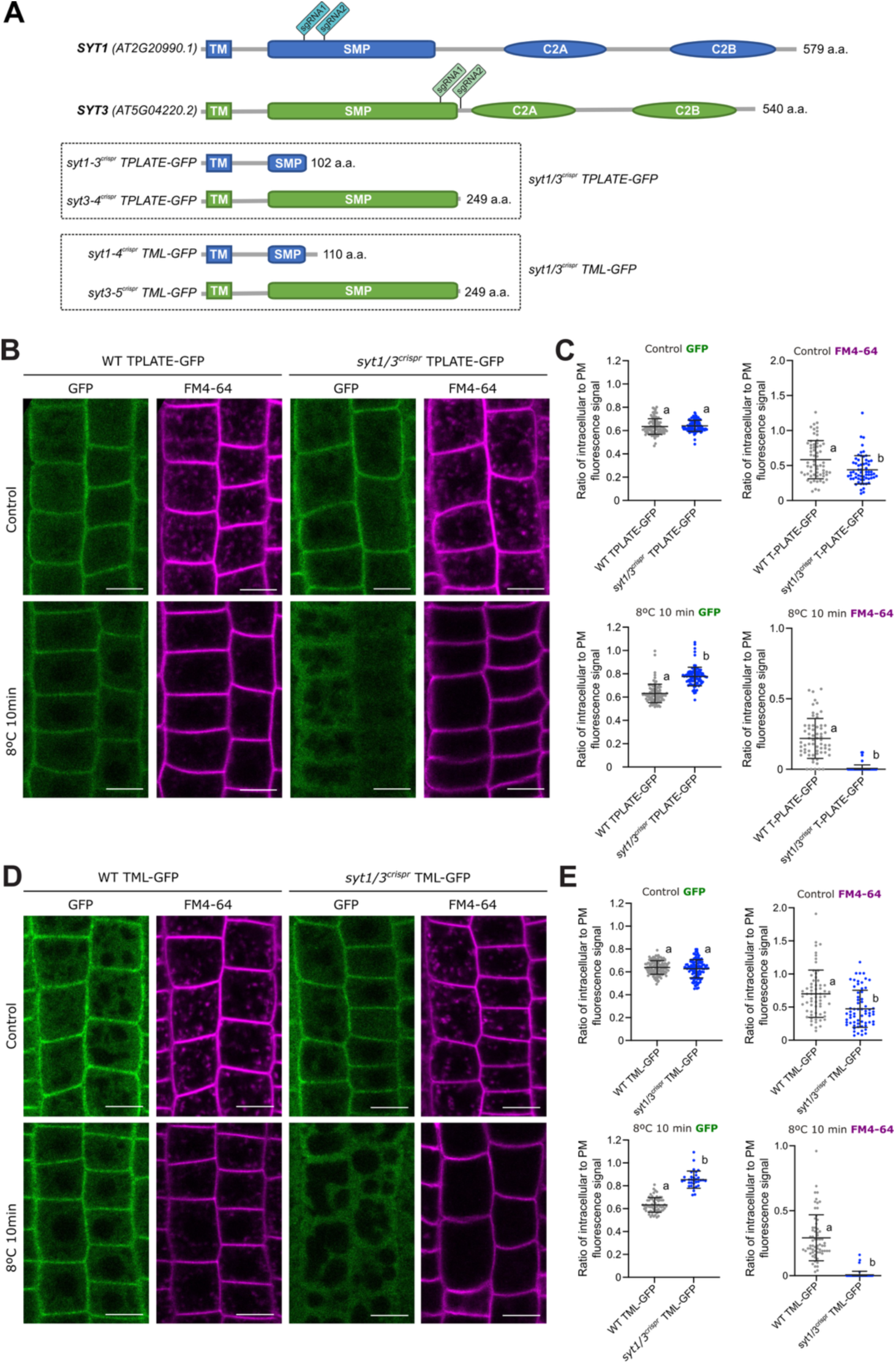
TPLATE and TML plasma membrane recruitment is abolished in the *syt1/3* double mutant under cold stress. (**A and B**) Representation of CRISPR sgRNA guides for *SYT1* and *SYT3* to generate mutant lines in the *tplate* and *tml-1* mutant backgrounds complemented respectively with TPLATE-GFP and TML-GFP. **(B and C)** Representative confocal images and quantification of the intracellular/PM ratio of TPLATE-GFP and FM4-64 internalization into endosomes under control and short-term cold treatment in 6-day old WT and *syt1/3* mutant root epidermal cells. **(C and D)** Representative confocal images and quantification of the intracellular/PM ratio of TML-GFP and FM4-64 internalization into endosomes under control and short-term cold treatment in 6-day old WT and *syt1/3* mutant root epidermal cells. **(C and E)** Individual dots represent cells. The lines represent the mean and the standard deviation (SD). Unpaired t-test with two-tailed *p*-value (*p*<0.05), n>100 cells for GFP and n>60 for FM4-64. For all assays, at least nine individual seedlings for each line were analyzed. Different letters indicate statistically significant differences. Scale bars, 10 µm.

Both *syt1/syt3* CRISPR mutant lines phenocopied the previously characterized *syt1/3* T-DNA double mutant, displaying clear defects in endocytosis. Under control conditions, FM4-64 internalization was reduced in both mutant backgrounds, revealing a basal endocytic defect, even though the intracellular-to-PM GFP fluorescence ratios of TPLATE-GFP and TML-GFP were comparable to those of their respective controls. Thus, loss of SYT1 and SYT3 does not detectably alter the steady-state PM association of these TPC markers under non-stress conditions but is sufficient to compromise endocytic uptake (Fig. 4b-e). Following cold treatment (8 °C, 10 min), the endocytic defect became much more pronounced: FM4-64 internalization was almost completely abolished in the *syt1/3crispr* lines, recapitulating the cold-induced phenotype observed in the *syt1/3* T-DNA double mutant (Fig. 1a-d). This strong inhibition of endocytosis coincided with impaired PM association of both TPC subunits, as shown by the loss of prominent peripheral GFP signal and increased intracellular-to-PM GFP fluorescence ratios relative to the corresponding controls (Fig. 4b-e). Together with the defective PM localization of the peripheral TPC subunit AtEH1/Pan1 (Fig. S4), these results indicate that SYT1 and SYT3 are required to maintain the PM association of both core and peripheral TPC components during cold stress. Because TPC inactivation blocks endocytosis (Wang et al. 2021), the loss of TPC membrane association provides a mechanistic explanation for the severe endocytic defect observed in cold-treated *syt1/3* mutants.

SYT1 and SYT3 play a role in clearing PM DAG generated by phospholipase C (PLC)-mediated phosphoinositide hydrolysis (Ruiz-Lopez et al. 2021), which is critical for the phosphoinositide cycle and replenishes PI4P and PI4,5P (Garcia-Hernandez et al. 2025; Ruiz-Lopez et al. 2021). Because TPC is recruited to the PM via charge-dependent protein-lipid interactions, short-term cold stress in the *syt1/3* mutant may profoundly affect the presence of negatively charged phosphoinositide isoforms in the PM. Moreover, treatment of Arabidopsis with the protonophore ES9 blocked clathrin-mediated endocytosis by immobilizing several core endocytic components (Dejonghe et al. 2016). In addition, ES9 reduced PI(4,5)P₂ at the PM, indicating that perturbing PM phosphoinositide abundance impairs endocytic machinery recruitment and dynamics (Dejonghe et al. 2016). Molecular dynamics simulations have also shown that anionic phospholipids recruit the hexameric core of TPC to the PM (Kraus et al. 2025). Therefore, reduced phosphoinositide replacement during cold stress, driven by a defective PI cycle in *syt1/3,* plausibly explains reduced TPC association with the PM and impaired endocytosis. To test our hypothesis, we used molecular dynamics simulations to determine whether increasing DAG at the expense of phosphoinositides affects TPC hexamer recruitment dynamics from the membrane. As previously described, the TPC hexamer interacted with PC, PE, PI4P, and PIP2 (Kraus et al. 2025) (Fig. 5a,b). In contrast, when phosphoinositides were depleted and DAG was increased, the TPC hexamer gradually lost its membrane localization, as indicated by the increasing distance between the protein center of mass and the center of mass of the phosphate headgroup atoms in the cytoplasmic leaflet (Fig. 5a,b). Collectively, these findings support a model in which cold-induced DAG accumulation in *syt1/3* disrupts the phosphoinositide-dependent electrostatic environment required for stable TPC association with the PM, providing a mechanistic explanation for impaired endocytosis under cold stress

**Figure 5.**
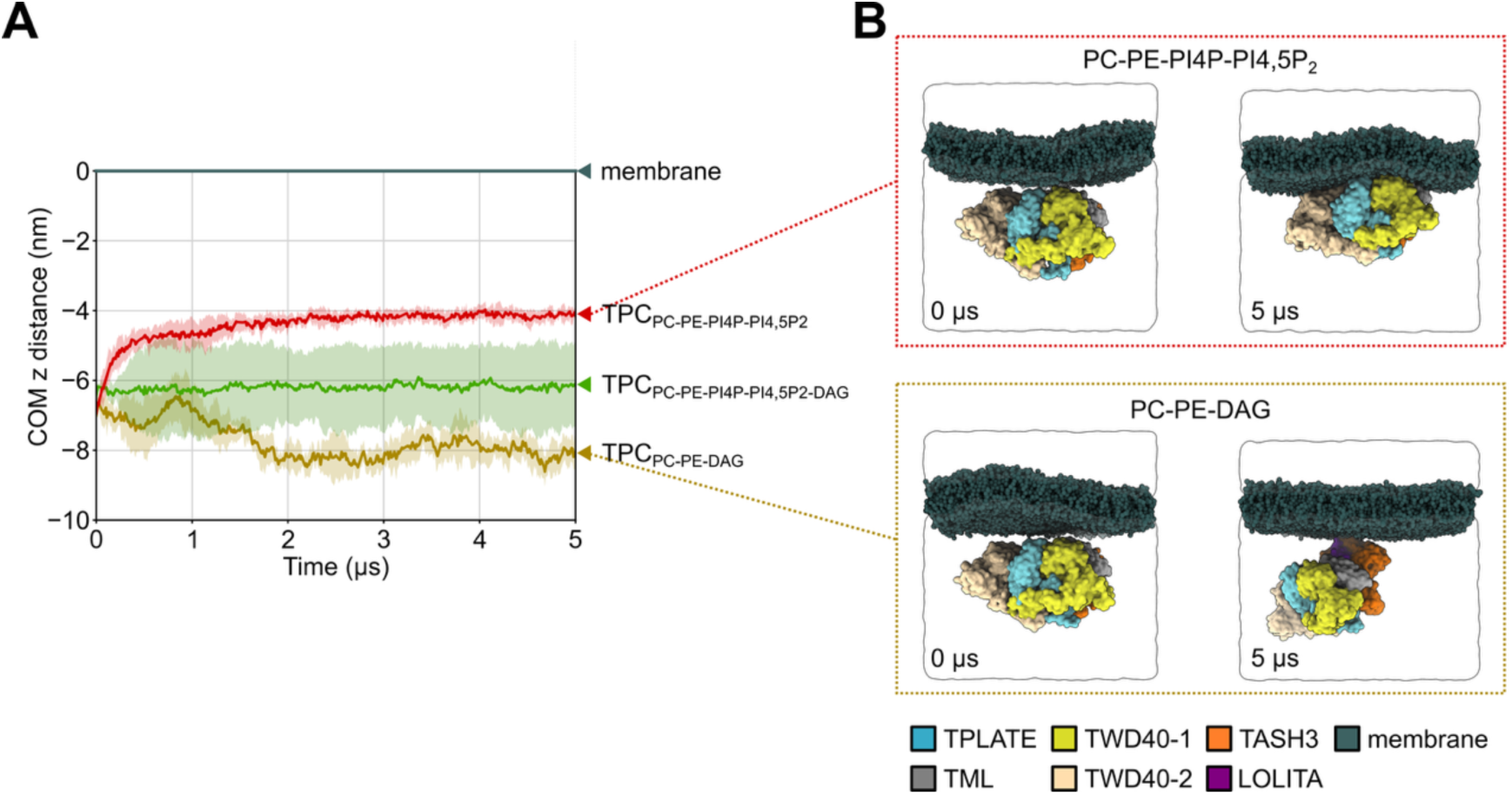
The TPLATE complex hexameric core loses membrane binding upon phosphoinositide depletion and increased DAG levels. (**A**) Z-distance between the center of mass (COM) of the PO₄ headgroups in the cytoplasmic leaflet and the hexameric part of the TPLATE complex (TPC) for different membrane compositions. TPC associated with the PC-PE-PI4P-PIP₂ membrane, showed mixed binding/not binding to the PC-PE-PI4P-PIP₂-DAG membrane, and exhibited little to no binding to the PC-PE-DAG membrane. Colours indicate different membrane compositions, transparent shading represents the standard error. (**B**) Representative simulation snapshots of the simulation box of TPC with PC-PE-PI4P-PIP₂ and PC-PE-DAG membranes at t = 0 and t = 5 µs.

## DISCUSSION

Our findings identify SYT1 and SYT3 as key determinants of endocytic activity during abiotic stress and link their functions at endoplasmic reticulum-plasma membrane (ER-PM) CS to the membrane association of the TPLATE complex (TPC). Here, we show that their loss severely compromises endocytosis during cold stress and reduces the PM association of both peripheral and core TPC subunits that, together with the molecular dynamics simulations, support a model in which SYT-dependent lipid homeostasis preserves the membrane environment required for endocytic machinery recruitment.

The contrasting phenotypes under control and stress conditions reveal a requirement for SYT1 and SYT3 that becomes particularly pronounced during cold exposure. Under control conditions, the *syt1/3* double mutant showed reduced endocytosis, whereas *syt1* alone showed no detectable defect. Cold treatment revealed a requirement for SYT1 and caused much stronger inhibition in *syt1/3*, indicating that SYT3 helps sustain endocytosis when SYT1 is absent. This functional redundancy is consistent with (1) their contributions to stress tolerance and PM lipid homeostasis, and (2) the high expression of *SYT1* compared with *SYT3* (Ruiz-Lopez et al. 2021). Recovery of FM4-64 uptake after return to control temperature indicates that the endocytosis defects are not due to irreversible cellular damage. The defective endocytosis of *syt1/syt3* under salt stress further extends the requirement for SYTs beyond cold, and is consistent with the original identification of *SYT1* in a role in NaCl tolerance (Schapire et al. 2008).

The loss of TPC membrane association provides a plausible explanation for the severe endocytic defect during cold stress. In cold-treated *syt1/3* mutants, the peripheral subunit AtEH1/Pan1 and the core subunits TPLATE and TML were depleted from the PM, which explains the inhibition of endocytic flux, as they are essential components of plant clathrin-mediated endocytosis (Gadeyne et al. 2014; Wang et al. 2021; Zhu et al. 2026). The effects on distinct parts of the complex support a broader failure to maintain TPC-membrane interactions. Moreover, independently generated CRISPR mutants reproduced the impaired FM4-64 uptake and altered localization of the core subunits, demonstrating the connection between SYT1 and SYT3 loss and these phenotypes. Although the experiments do not establish whether TPC displacement precedes endocytic inhibition, the requirement for TPC makes its impaired PM association a likely contributor. Interestingly, control conditions also show that reduced FM4-64 uptake occurred even though TPLATE and TML remained at the PM. Thus, the basal endocytic phenotype cannot be explained by an obvious redistribution defect of TPC, as observed during cold stress. Loss of SYT1 could affect membrane deformation, endocytic-complex dynamics, or subsequent trafficking steps without producing a detectable change in steady-state localization. This distinction suggests that membrane association alone does not fully capture endocytic competence. Cold exposure may intensify an existing membrane defect to the point that TPC can no longer maintain stable association.

The lipid-binding properties of TPC provide a possible mechanistic framework for interpreting these observations. Anionic phospholipids contribute to the electrostatic identity of the plant PM, with PI4P playing a major role (Simon et al. 2016), and lipid interactions support the membrane association of TPC components (Yperman et al. 2021a). We have previously shown that during cold stress, the *syt1/syt3* mutant accumulates excessive DAG in the PM (Ruiz-Lopez et al. 2021). Because DAG influences membrane packing and curvature, its accumulation will affect the physical environment in which endocytic complexes assemble (Campomanes et al. 2019). Moreover, work in animal cells has shown that local DAG gradients can guide the movement of clathrin-coated pits and facilitate exocytosis-coupled endocytosis, supporting the idea that DAG accumulation can directly influence the membrane environment required for endocytic dynamics (Yuan et al. 2015). Perturbations in phosphoinositide metabolism could also alter the availability of negatively charged lipids required for TPC binding. This is supported by molecular dynamics simulations in which phosphoinositide depletion combined with increased DAG strongly affects the association of the TPC hexameric core with the lipid bilayer.

The effects on BRI1 distribution in *syt1/syt3* mutant extend the significance of these findings to PM receptor trafficking and signaling. During cold stress, the intracellular-to-PM BRI1-GFP ratio increased in wild type but not in *syt1*, consistent with reduced receptor internalization in the mutant. In parallel, *syt1/3* seedlings showed increased BES1 dephosphorylation and higher expression of BR-responsive genes, consistent with the relationship between BRI1 trafficking and BR signaling (Irani et al. 2012; Martins et al. 2015; Nolan et al. 2020). These data suggest that SYT-dependent membrane maintenance helps preserve receptor trafficking during stress, with potential consequences for coordinating growth and environmental responses.

The association between SYT1 and VAP27-1 further connects ER-PM CS with endocytic regulation. VAP27 proteins have previously been implicated in clathrin dynamics and endocytic trafficking (Stefano et al. 2018). Co-immunoprecipitation and proteomic analysis support an association of VAP27 with SYT1, whereas the absence of detectable FRET does not establish a direct interaction. These findings support an indirect association within a larger protein assembly or interactions not detected by FRET. The genetic results also indicate a condition-dependent relationship: the *syt1/vap27-1/3* triple mutant resembled *vap27-1/3* under control conditions and *syt1* during cold stress. This non-additivity is consistent with overlapping contributions to endocytosis, but does not establish a linear pathway or an exclusive division between basal and stress functions. Together, the data suggest that distinct ER-PM tethering systems contribute differently, one through direct interaction with the endocytic machinery (VAP27s) and the others through controlling proper lipid homeostasis at the PM (SYTs). The transcriptional analysis provides additional evidence that SYT loss perturbs cellular homeostasis. Enrichment of wound– and jasmonate-associated responses under control conditions is consistent with a disturbance that becomes more pronounced after cold stress. The larger transcriptional differences between cold-treated *syt1/3* and WT, including enrichment of lipid-related responses, support a more severe trafficking phenotype. Altered lipid organization, receptor trafficking, and stress signaling may contribute jointly to the transcriptional response. This reciprocal relationship is relevant because endocytosis depends on membrane properties and helps maintain membrane composition.

Overall, these results position SYT1 and SYT3 as functionally linking membrane homeostasis and endocytic trafficking by maintaining TPC membrane association during cold stress, and they support a lipid-based mechanism for this requirement. By preserving a membrane environment compatible with endocytic activity, ER-PM contact sites may help sustain membrane turnover and receptor regulation during environmental challenges.

## MATERIAL AND METHODS

### Plant material and growth conditions

*Arabidopsis thaliana* (Arabidopsis) and *Nicotiana benthamiana* plants were used in this study. All Arabidopsis plants were of Columbia-0 (Col-0) ecotype except for the *vap27-3* mutant (Sail_355_E10) which is in Col-3 background. The *Arabidopsis* single mutant lines used in this study were *syt1-2* (AT2G20990) SAIL_775_A08 (Pérez-Sancho et al. 2015) and *syt3-1* (AT5G04220) SALK_037933 (Ruiz-Lopez et al. 2021). Double null mutant lines for *syt1/syt3* and *vap27-1vap27-3* (*vap27-1/3*) (WiscDsLoxHs096_06H, Col-0 background and Sail_355_E10, Col-3 background for *vap27-1* and *vap27-3* respectively) were also previously reported (Ruiz-Lopez et al. 2021; Stefano et al. 2018), while triple mutant *syt1/vap27-1-1/3* was obtained by crossing *syt1* and *vap27-1/3*. The identification of homozygous plants carrying the T-DNA insert in each mutant was confirmed through diagnostic PCR. Allele-specific primers listed in Supp. Data 1a, were employed for this purpose. The transgenic lines used were *SYT1:SYT1-GFP* in *syt1* (Pérez-Sancho et al. 2015), *SYT5:SYT5-GFP* in *syt5* (García-Hernández et al. 2025), *35S:GFP* (Wang et al. 2019), *BRI:BRI1-GFP* (Geldner et al. 2007), *VAP27:VAP27-1-GFP* (Wang et al. 2016) *TPLATE-GFP* (pLAT52-TPLATE-GFP in *tplate*) (Van Damme et al. 2006), *TML-GFP* (pTML-TML-GFP in *tml-1*) (Gadeyne et al. 2014). To generate the transgenic line expressing BRI1 fused to GFP under the control of the endogenous BRI1 promoter, *syt1* homozygotes mutant plants were genetically crossed with *BRI1:BRI1-GFP* plants (Geldner et al. 2007) and were selected with fluorescence for BRI1-GFP and further identify by diagnostic PCR for *syt1*. Homozygous *BRI1:BRI1-GFP* in *syt1 (syt1/BRI1-GFP)* were confirmed by analysis the GFP fluorescence of the progeny. The *syt1/3* stable transgenic doble mutant lines in the TML-GFP and T-PLATE-GFP background was generated as described in the “Plasmid constructs” and “Generation of transgenic plants” sections. When required, seedlings were transferred to soil after 7 days of in vitro growth and watered every 2 days. In soil, plants were grown in a mixture of organic substrate and vermiculite (4:1, v/v) at 22°C, with a long-day (16/8 h day/light cycles) photoperiod (approximately 120 μmol photon m^−2^ s^−1^).

### Plasmid constructs

For generation of *syt1/3* doble mutant crispr lines in the TML-GFP and T-PLATE-GFP background, two CRISPR guides directed against *SYT1* and two CRISPR guides directed against *SYT3* were designed using the CHOPCHOP tool (Labun et al. 2019) (https://chopchop.cbu.uib.no/). For the guides sequences, see Supplementary Data 1a. The plasmids for plant transformation were prepared using the CRISPR-Cas9 cloning system and protocol developed in Stuttmann et. Al. (Stuttmann et al. 2021). For plasmid list and description, see Supplementary Data 1b. The following plasmid (Supp. Data 1b) constructs used in this study were previously described: *35S:SYT1-GFP* and *UBQ10:SYT1-Cherry* (Ruiz-Lopez et al. 2021). *UBQ10:VAP27-1-cherry* was a kindly provided by Stephen Vanneste. *UBQ10:Cherry* was generated by LR multiple gateway reaction using the the *pENTR* vectors *pEN L4-pUBQ10-R1, pEN pEN L1-mCherry-L2 and pDONRP2R-P3 (Mock), with the pDEST vector pH7m34GW*.

### Generation of transgenic plants

The CRISPR/Cas9 expression constructs used to generate the *syt1/3* crispr TML-GFP and T-PLATE-GFP lines were transformed into *Agrobacterium tumefaciens* strain GVG3101::pMP90 through electroporation and confirmed by diagnostic PCR. Then, Arabidopsis plants were transformed by foral dip (Clough and Bent 1998) to generate stable transgenic mutant plants. The selection process was executed using the identification of red fluorescent seeds, since transgenic plants carried the FastRed selection system. Immunoblot analysis of plants from red fluorescent seeds using the anti-SYT1 antibody (see immunoblot section) were performed to select loss-of-function mutants for *SYT1*. Next, the molecular nature of the *syt1* mutation was investigated by sanger sequencing. The lines that were confirmed as *syt1* knock-out mutant by western blot and sequencing, were further analysis by sequencing of the *SYT3* CRISPR/CAS9 guides target region. Plants presenting knock-out alleles for *SYT3* were identified and T3 and T4 homozygous *syt1/3* crispr lines TML-GFP and T-PLATE-GFP lines were selected and used in this study.

### Cold treatment and sample preparation for microscopy live-cell imaging

Six-days old seedlings grown at 22 °C with 16/8 h day/light cycles in ½ agar-solidified MS medium supplemented with 1.5% sucrose were treated with or without short-term cold treatment (8 °C, 10 min) on ¼ MS liquid medium (supplemented with 1.5% sucrose) and then were mounted on microscope slides and cover slips and imaged with Zeiss confocal microscope from 5 to 10 minutes after mounting (Fig. S1). A drop of ddH_2_O was added to the microcopy slide to accommodate seedlings and an agarose pad prepared with 0.8% agarose in ¼ MS medium (supplemented with 1.5% sucrose) was used for mount seedling, immobilizes it for high-resolution imaging.

### FM4-64 uptake and imaging in cold assays

For FM4-64 uptake analysis (Fig. S1), seedlings were incubated immerse in ¼ MS medium (supplemented with 1.5% sucrose) containing 2µM FM4-64 (Invitrogen) for 10 min, briefly washed in the ¼ MS medium (supplemented with 1.5% sucrose) to remove excess of FM4-64 and observed from 5 to 10 min after 8 °C by confocal imaging. During the staining, the staining solution was protected from light. Images were taken using a laser scanning confocal microscopy (LSCM, Zeiss 880). Experiments were performed in three independent days and for each experiment, at least four independent seedlings were performed. For FM4-64, samples were excited at 561 nm and 488 nm respectively and detected at 620-700 nm (FM4-64).

### BRI1-GFP imaging in cold assays

For BIR1-GFP imaging, 6-days old seedling were grown and treated as described in the “Cold treatment and sample preparation for microscopy live-cell imaging” section. Images were taken using a laser scanning confocal microscopy (LSCM, Zeiss 880). Experiments were performed in three independent days and for each experiment, at least four independent seedlings were performed. For GFP signal detection, samples were exited at 488 nm and detected at 500-550 nm.

### FM4-64 uptake and imaging in salt assays

For salt treatment, seedlings were pretreated immerse in ¼ MS medium (supplemented with 1.5% sucrose) with 100 mM NaCl for 50 min and then further incubated for 10 min with 2µM FM4-64 (Invitrogen), briefly washed in the ¼ MS medium (supplemented with 1.5% sucrose) to remove excess of FM4-64 and observed from 5 to 10 min after 8 °C by confocal imaging. As described above, images were taken using a laser scanning confocal microscopy (LSCM, Zeiss 880) and experiments were performed in three independent days and for each experiment, at least four independent seedlings were performed. For FM4-64, samples were excited at 561 nm and 488 nm respectively and detected at 620-700 nm.

### BFA treatment and imaging in cold assays

For BFA treatment, 6-days old Arabidopsis seedlings were incubated immerse in liquid ¼ MS medium (supplemented with 1.5% sucrose) containing BFA (50 µM) inhibitors for 50 min and then further incubated for 10 min with 2µM FM4-64 (Invitrogen), briefly washed in the ¼ MS medium (supplemented with 1.5% sucrose) to remove excess of FM4-64 and observed from 5 to 10 min after 8 °C by confocal imaging as described above.

### Combined imaging of GFP and FM4-64

Arabidopsis root images for quantification of intracellular relative to PM fluorescence signal intensity for TPLATE-GFP and TML-GFP in double mutant *syt1/3* background were obtained using a confocal Zeiss LSM 880 confocal microscope and using the Zen Black Software (Zeiss). Images were acquired with a ×40 water-immersion corrected objective, frame scan mode (1024 × 1024) and four-line bidirectional signal averaging and with a ×2 digital zoom. A single excitation (488 nm laser) line and emission windows ranging between 500 and 541 nm (GaAsI detector, chanel 2) for GFP; and 612-759 nm for FM4-64 (T13320 ThermoFisher) were used. Experiments were performed in three independent days and for each experiment, at least four independent seedlings were analyzed.

### Quantifications of intracellular relative to PM fluorescence signal ratios

Quantification of intracellular relative to PM fluorescence signal ratios for both FM4-64 and BRI1-GFP was performed as previously described (Rigal et al. 2015) with some modifications, using the Fiji software package (ImageJ v2.16.0/1.54p; (Schindelin et al. 2012; Schindelin et al. 2015). Briefly, to eliminate background noise without removing signal from endocytic structures, a 20% intensity threshold was established and uniformly applied to all images prior to quantification in Fiji. Two distinct regions of interest (ROIs) were defined per cell: a polygonal ROI with a line thickness of 5 pixels tracing the PM, and a second ROI encompassing the entire intracellular area enclosed by the PM polygon (intracellular). After applying the 20% threshold, cells without detectable endocytic structures had an intracellular signal value of zero. The mean intensity of the 5% brightest pixels within the intracellular ROI was divided by the mean intensity of the 5% brightest pixels within the PM ROI. Only confocal images without saturated pixels were included in the analysis.

Quantifications of intracellular relative to PM GFP fluorescence signal ratios in the experiments with TPLATE-GFP and TML-GFP (combined imaging of GFP and FM4-64), were performed with a custom-made Fiji-based script previously described (Grones et al. 2022) and available at https://github.com/pegro-psb/Cyto-PM-signal-quantification. With this script we used the FM4-64 staining in the red channel as a mask to allow automatic detection of PM and cytoplasm regions. Then, GFP and FM4-64 images were merged in Fiji. Cells were manually annotated and stored in the Fiji ROI Manager. The script automatically measures the mean intensity of the top 5% brightest pixels of the channel 2 (GFP) in the detected PM and cytoplasm ROI of each annotated cell based on the FM4-64 signal in channel 1 (FM4-64). We determined the intracellular-to-PM fluorescence ratio by dividing the mean intensity of the top 5% brightest intracellular pixels by the mean intensity of the top 5% brightest PM pixels.

### Phosphorylation Status of BES1

Six-days old seedlings grown at 22 °C with 16/8 h day/light cycles in ½ agar-solidified MS medium supplemented with 1.5% sucrose were treated with short-term cold treatment (8 °C) on ¼ MS liquid medium (supplemented with 1.5% sucrose) and frozen in liquid nitrogen after 0, 0.5, 1 and 2 hours of treatment. Approximately 200 mg of ground tissue per sample was used for total protein extraction. Denatured protein extracts were obtained by homogenizing and incubating plant material in 200 mL of 2X Laemmli buffer (125 mM Tris-HCl, pH 6.8, 4% [w/v] SDS, 20% [v/v] glycerol, 2% [v/v] b-mercaptoethanol, and 0.01% [w/v] bromophenol blue) for 5 min at 95°C, centrifuged (for 5 min at 20,000g) at room temperature. The total proteins from supernatant were separated in a 10% SDS-PAGE gel and analyzed using an anti-BES1 antibody (dilution 1:500) (Yu et al. 2011) as described under “Immunoblot” section.

### Confocal microscopy and FRET in *Nicotiana benthamiana*

*N. benthamiana* plants were grown in a greenhouse at 22 °C under 16/8 light/dark conditions. Transient expression was performed by leaf infiltration of *Agrobacterium tumefaciens* (GV3101::pMP90) carrying the different constructs, together with the Agrobacterium expressing the p19 silencing suppressor. Similar optical density of Agrobacterium strains expressing the constructs were used during co-expression. Cells were then harvested by centrifugation, pellets were resuspended in agroinfiltration solution (10 nM MES, pH 5.6, 10 mM MgCl_2_ and 1 nM acetosyringone) and incubated for 2 h in dark conditions at room temperature. Three to four-week-old *N. benthamiana* leaves were infiltrated with Agrobacterium into the abaxial side of the leaf lamina. Transiently transformed *N. benthamiana* were imaged from two to three days after infiltration using a laser scanning confocal microscopy (LSCM, Zeiss 880). For each experiment, at least three independent infiltrations were performed. Images were taken in multi-track mode with frame switching when multifluorescence was used. For GFP/Cherry combination, samples were excited at 488 and 587 nm and detected at 500-550 and 580-630 nm, respectively.

### RNA extraction and RNAseq analysis

RNA extraction was performed using a RNeasy plant mini kit (Qiagen, http://www.qiagen.com/). Reverse transcription experiments were performed using a Superscript III first-strand synthesis kit (Invitrogen, http://www.invitrogen.com/). For RNAseq analysis, 6-days old seedlings treated with or without short-term cold (8 °C, 10 min) treatment were powdered in liquid N_2_ and total RNA was isolated using the E.Z.N.A.® Plant RNA Kit (Omega Bio-Tek) and the clustering of the index-coded samples was performed on a cBot Cluster Generation System using TruSeq PE Cluster Kit v3-cBot-HS (Illumina) according to the manufacturer’s instructions. After cluster generation, the library preparations were sequenced on an Illumina Novaseq platform, and 150 bp paired-end reads were generated at the Novogene (UK) Company Ltd. At last, RNA integrity was assessed using the RNA Nano 6000 Assay Kit of the Bioanalyzer 2100 system (Agilent Technologies, CA, USA). The reads were quality filtered and trimmed using Trimmomatic version 0.36 (Bolger et al. 2014) with default paired-end mode options. The resulting reads were then aligned to the TAIR10 version of the *Arabidopsis thaliana* genome sequence (https://www.arabidopsis.org/) using Hisat2 version 2.1.0 (Kim et al. 2015). These read alignments (in BAM format) were used for transcript quantification with the cuffdiff program of the Cufflinks version 2.2.1 package (Trapnell et al. 2013). The resulting read alignments were visualized and clustered using Tablet software (Milne et al. 2013) and CummeRbund R package version 2.23.0 (Goff et al. 2013). To determine DEGs, a False Discovery Rate (q-value) cutoff of ≤ 0,05 and a Log2 Fold Change of 1 was set. DEGs (input list) were subjected to singular enrichment analysis for the identification of overrepresented GO terms using Panther database available at: http://go.pantherdb.org/tools/compareToRefList.jsp (Mi et al. 2019a; Thomas et al. 2022; Mi et al. 2019b) with the default options (Released 20231017, GO Ontology database DOI: 10.5281/zenodo.8436609 Released 2023-10-09). The Fisher’s exact test with the FDR correction as calculated by the Benjamini-Hochberg procedure was used and only results with FDR< 0.05 were selected for further analysis. Bubble graph was plotted with GraphPad Prism 10. Arabidopsis RNAseq transcriptome proceeded data is available in Supp. Data 2.

### Immunofluorescence

For the immunolocalization four days old seedlings treated with a short-term cold treatment (8 °C, 10 min). Immediately after the treatment approximately 20-30 seedlings per sample were transferred into 4% PFA (in PBS buffer) and fixated under vacuum for 60 min. The fixated seedlings were transferred into InsituPro VSI roboto (Invavis) and immunolocalization was performed as described in (Sauer and Friml 2010). Briefly, seedlings were incubated in 2% Driselase in PBS (Japan) for 30 min, in 1,5% NP-40 in PBS (Sigma) for 30 min and in 2% BSA in PBS (Sigma) for 30 min. Afterwards the samples were incubated for 4 hours with primary antibody α-AtEH1/Pan1 (rabbit – (Grones et al. 2022)) and secondary antibody α-Rabbit-Alexa Fluor-488 (Invitrogen, A11034) were used in dilution 1:600. After the immunolocalization, seedlings were kept in the DAPCO antifading solution (Sigma). Samples were visualized using a confocal microscope Leica Stellaris 5 equipped with 20x objective (NA 0.75) and white light lase (WLL) with excitation of 488 nm and emission range of 500-585 nm.

### Co-Immunoprecipitation

Arabidopsis seedlings were grown for 6 days on ¼ agar-solidified MS medium supplemented with 1.5 % agarose at 22°C with 16/8 light/dark photoperiods. Seedlings were harvested, flash-frozen and ground in liquid nitrogen. Proteins were extracted in a 1:1 ratio (50 mM Tris-HCl, pH 7.5; 150 mM NaCl; 10% glycerol; 10 mMEDTA, pH 8; 1 mM NaF; 1 mM Na_2_MoO_4_.2H_2_O; 10 mM DTT; 0.5 mM PMSF; 1% (v/v) P9599 protease inhibitor cocktail; Nonidet P-40 0.5% (v/v.). Protein extracts were incubated for 20 min at 4°C on a rotation wheel and cleared by centrifugation at 20,000g for 30 min. The protein content of the supernatant was incubated with 20 µL GFP trap agarose beads (Chromoteck) for 1 hour at 4°C on a rotation wheel. Beads were washed three times with the lysis buffer free of detergent. Then, samples were analyzed by loading on 12% gels after addition of 4X Laemmli sample buffer and boiling at 70°C for 45 min and gels were transferred to PVDF membranes using the Trans-Blot Turbo system (Bio-Rad) and analysis as described in the “Immunoblot analysis” section.

### Immunoblot analysis

Proteins separated by SDS-PAGE were blotted using Trans-blot Turbo Transfer System (Bio-Rad) onto polyvinylidene difluoride (PVDF) membranes (Immobilon-P, Millipore) following the instructions by the manufacturer. PVDF membranes, containing electroblotted proteins, were then incubated with the appropriate primary antibody followed by the appropriate secondary peroxidase-conjugated antibody. The following primary antibodies were used for detection of epitope-tagged proteins: rabbit polyclonal anti-BES1 antibody (dilution 1:500; (Yu et al. 2011)), mouse monoclonal anti-GFP clone B-2 (1:1000; catalog no. sc-9996, Santa Cruz Biotechnology), and rabbit polyclonal anti-SYT1 antibody (1:5000). The secondary antibodies used in this study were anti-mouse immunoglobulin G (IgG) whole-molecule peroxidase (1:80,000; catalog no. A9044, Sigma-Aldrich) and anti-rabbit IgG whole-molecule peroxidase (1:80,000; catalog no. A0545, Sigma-Aldrich). Proteins and epitope-tagged proteins on immunoblots were detected using the Clarity ECL Western Blotting Substrate or SuperSignal West Femto Maximum Sensitivity Substrate according to the manufacturer’s instructions, and images of different time exposures were acquired using the Chemidoc XRS1 or MP System (Bio-Rad). Only images with no saturated pixels were used for protein quantification. To confirm equal loading of the different samples in each experimente, immunoblotted PVDF membranes were stained with Coomassie brilliant blue R-250 or the stain-free imaging (BioRad) of the PVDF membrane blot was used.

### Molecular dynamics simulations

Coarse-grained molecular dynamics simulations were performed using GROMACS software (Abraham et al. 2015) and the MARTINI 2.2 coarse-grained force field (de Jong et al. 2013). The previously generated MARTINI 2.2 coarse-grained model of the TPC hexamer was used as the starting structure (Kraus et al. 2025).

Three 25 × 25 nm membrane compositions were generated using the insane.py script (Wassenaar et al. 2015). The PC-PE-PI4P-PI(4,5)P2 membrane consisted of 50% PC and 50% PE in the upper leaflet and 46% PC, 46% PE, 5% PI4P, and 3% PI(4,5)P2 in the lower leaflet. The PC-PE-PI4P-PI(4,5)P2-DAG membrane consisted of 50% PC and 50% PE in the upper leaflet and 46% PC, 46% PE, 1% PI4P, 0.6% PI(4,5)P2, and 6.4% DAG in the lower leaflet. The PC-PE-DAG membrane consisted of 50% PC and 50% PE in the upper leaflet and 46% PC, 46% PE, and 8% DAG in the lower leaflet.

TPC was positioned in proximity to the pre-equilibrated membrane patch and inserted into a 25 × 25 × 27.5 nm simulation box. Subsequently, the systems were solvated with 0.15 M NaCl in water and neutralized. The systems were energy-minimized using the steepest-descent algorithm for a maximum of 5,000 steps and equilibrated for 4,750 ps. Position restraints on the protein backbone and side chains were progressively released during equilibration. For each membrane composition, five independent 5 μs long production runs, starting from different initial velocities, were performed with a 20-fs time step in the NPT ensemble. Pressure was set to 1 bar and maintained using the Parrinello–Rahman barostat, with a coupling constant of 12.0 ps and a compressibility of 3.4 × 10−4. Temperature was set to 323 K and maintained using the v-rescale thermostat with a coupling constant of 1.0 ps. Bond lengths were constrained using the LINCS algorithm. Van der Waals and Coulomb cut-offs were set to 1.1 nm.

Z-distance between protein COM and COM of the lower leaflet’s phosphate headgroups was calculated using GROMACS in-build tool *gmx traj*. Simulation snapshots were generated using UCSF ChimeraX software (Pettersen et al. 2021; Meng et al. 2023).

### Statistical analysis

For statistical analyses, the One or Two-way analysis of variance with Tukey’s multiple comparison test, Two-way ANOVA with Šidák’s multiple comparison test, and Unpaired t-test with two-tailed *p*-value was performed using GraphPad Prism version 5 and 9 as described in figures legends. A *P* value < 0.05 was considered as statistically significant. Statistical methods used to calculate *P* values are described in the figure legends.

## AUTHOR CONTRIBUTIONS

J.A.A.-M., Á.D.E., J.P.-S., M.N., P.G., R.P., D.V.D., V.A.-S. and M.A.B. conceived and designed the research.

J.A.A.-M., Á.D.E., V.A.-S. performed most of the experiments. M.N. and R.P. performed the Molecular dynamics simulations experiments. G.W. and P.G. performed Immunofluorescence experiments. J.M.-C. extracted RNA for RNA-seq, assembled the first transcriptome, and analyzed the data. J.M.-C., J.A.A.-M., V.A.-S. and M.A.B. curated and analyzed transcriptomic data. A.D.M. and S.V. helped with some experiments.

J.A.A.-M., Á.D.E., J.P.-S., M.N., P.G., R.P., L.R., D.V.D., V.A.-S. and M.A.B. analyzed and discussed the data.

J.A.A.-M., M.N., R.P., D.V.D., V.A.-S. and M.A.B. wrote the manuscript with the contribution of all other authors.

All the authors have read, revised and approved the paper.

## ACKNOWLEDGMENTS

We would like to thank Federica Brandizzi at Michigan State University for kindly provided *vap27-1/3* Arabidopsis mutant line. We thank Pengwei Wang from the Huazhong Agricultural University (Wuhan, China) for kindly provided Arabidopsis line expressing VAP27-1-GFP, and Yanhai Yin Iowa State University for generously providing the anti-BES1 antibody.

We would also like to thank to the computational resources used for molecular dynamics simulations were provided by the e-INFRA CZ project (ID: 90254), supported by the Ministry of Education, Youth and Sports of the Czech Republic.

## FUNDING

Spanish “Ministerio de Ciencia Innovación y Universidades” (MICIU), PID2023-147983OB-I00 (M.A.B and L.R.).

“Programa Emergia 2023” grant (DGP_EMEC_2023_00375) from the “Consejería de Universidad, Investigación e Innovación de la Junta de Andalucía” (V.A-S.).

Marie Skłodowska-Curie actions, The European Research Executive Agency, H2020-MSCA-IF-2019 grant no. 893075-PLISUS (J.A.A-M.).

Spanish “Ministerio de Ciencia Innovación y Universidades” (MICIU), PID2022-138838OA-I00 (J.A.A-M.).

Spanish “Ministerio de Ciencia Innovación y Universidades” (MICIU) and co-financed by MICIU/AEI and FSE+, RYC2024-049082-I (J.P-S.).

Ministry of Education, Youth and Sports of the Czech Republic, GA UK number 154324, e-INFRA CZ project ID: 90254 (M.N.).

Funding for open access charge: Universidad de Málaga/CBUA.

## SUPPLEMENTAL DATA

**Supplemental Data 1.** Primers (a) and plasmids (b) used and generated in this work.

**Supplemental Data 2.** Arabidopsis RNAseq transcriptome of 6-days old seedlings WT (Col-0) and *syt1/3* in control and cold stress (8 °C, 10 min) treatment.

## SUPPLEMENTAL FIGURES

**Figure S1.**
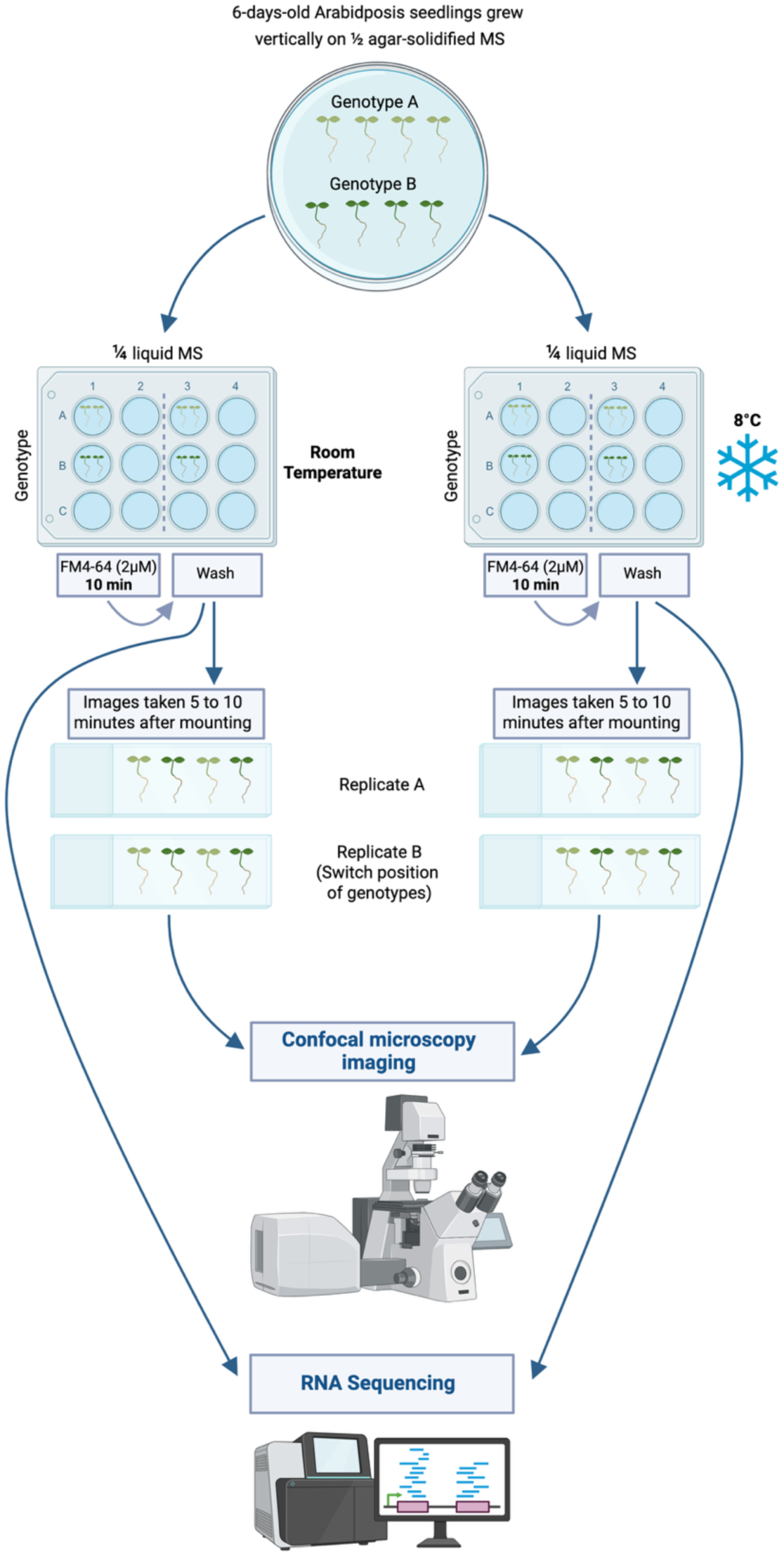
Schematic representation of the workflow used in the analysis. Six-day-old seedlings treated with or without short-term cold treatment (8 °C, 10 min immersed in ¼ MS liquid medium followed by recovery at room temperature for 5 min) were processed for confocal microscopy and RNA sequencing. For FM4-64 uptake, seedlings were incubated with ¼ MS medium containing 2µM FM4-64 (Invitrogen) for 10 min and observed after 5 to 10 min. During the staining, the FM4-64 staining solution was protected from light. Created in https://BioRender.com

**Figure S2.**
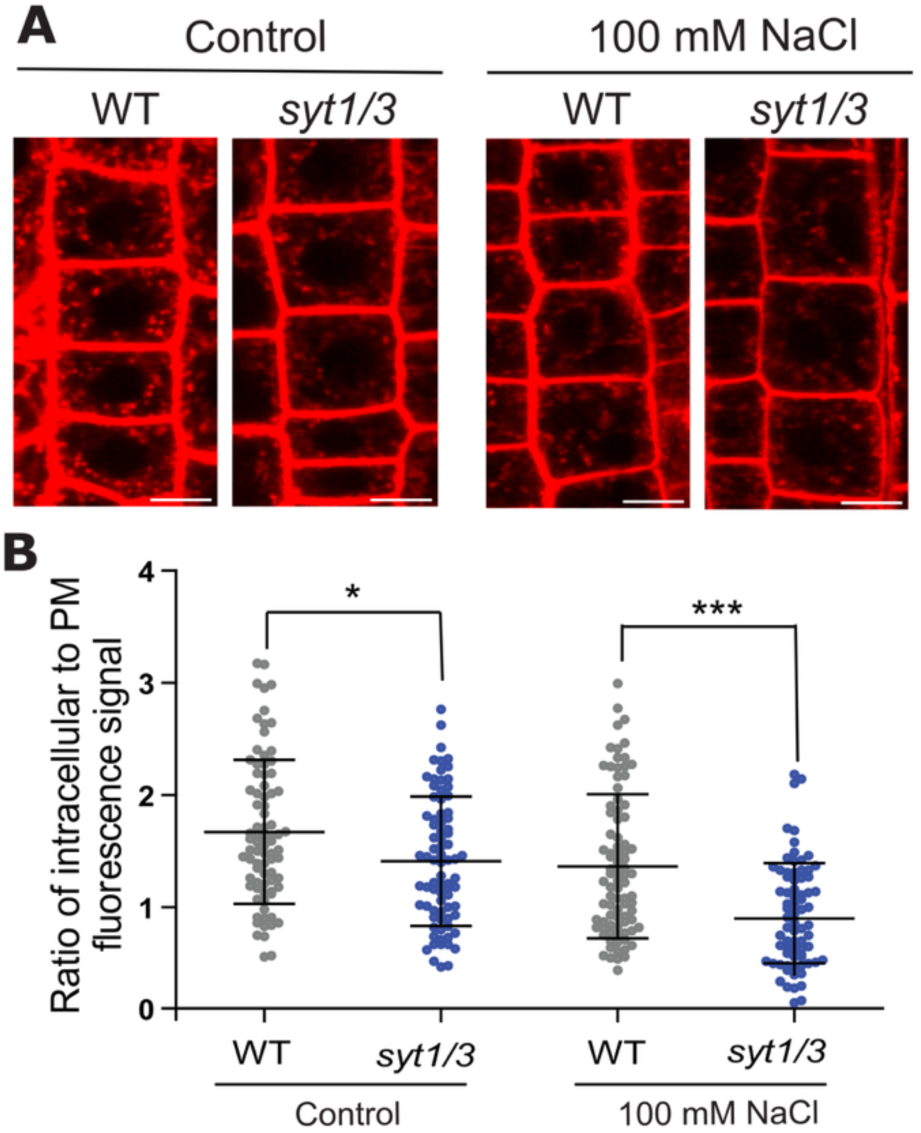
*syt1/3* mutant impairs endocytosis under salt stress. **(A)** Six-day-old seedlings treated without (control) or with salt (100mM NaCl) for 50 min on ¼ MS liquid medium were incubated with 2µM FM4 64 (¼ MS medium) for 10 min and visualized by confocal imaging. **(B)** Quantification of the fluorescence ratio between intracellular endosomal and PM signal of the images in panel A. Individual dots represent cells. The lines represent the mean and the standard deviation (SD). Scale bar, 5 µm. One way-ANOVA with Tukey’s multiple comparison test and *p*<0.05, n>75 cells.

**Figure S3.**
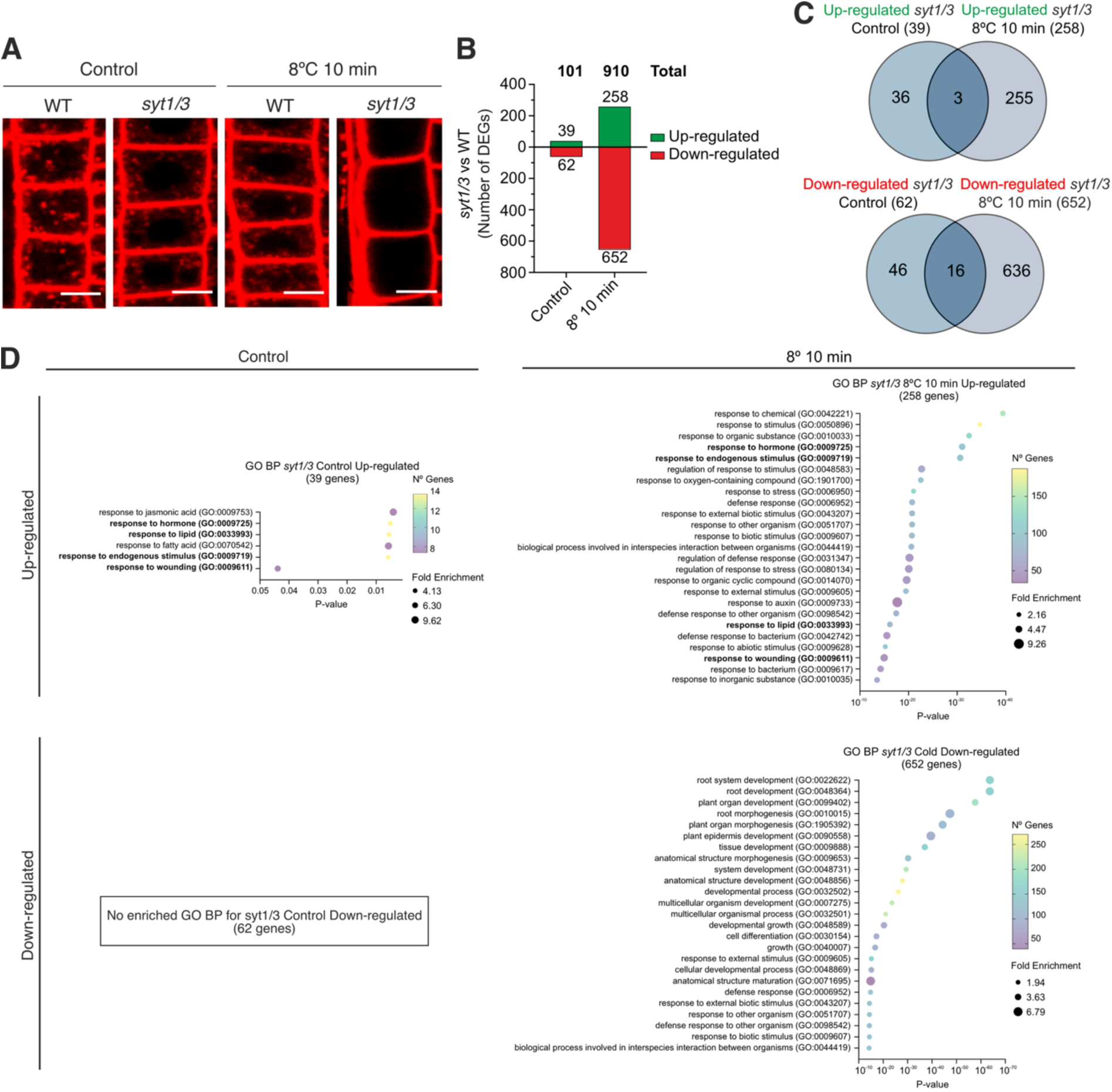
RNAseq analysis of *syt1/3* under control and cold stress. **(A)** Representative confocal images and quantification of FM4-64 internalization under control and short-term cold treatment in 6-day old WT and *syt1/3* mutants root epidermal cells. (**B**) Number of differentially expressed genes (DEGs) between *syt1/3* vs WT comparisons both in control and cold conditions. Green bars represent induced genes vs WT. Red bars represent repressed genes vs WT. **(C)** DEGs overlap between *syt1/3* control vs WT control and *syt1/3* cold vs WT cold. **(D)** Gene ontology for “Biological Process” category for *syt1/3* vs WT control and cold DEGs using the Panther online tool with default options. GO categories that are enriched in both, control and cold for *syt1/3* induced genes are in bold.

**Figure S4.**
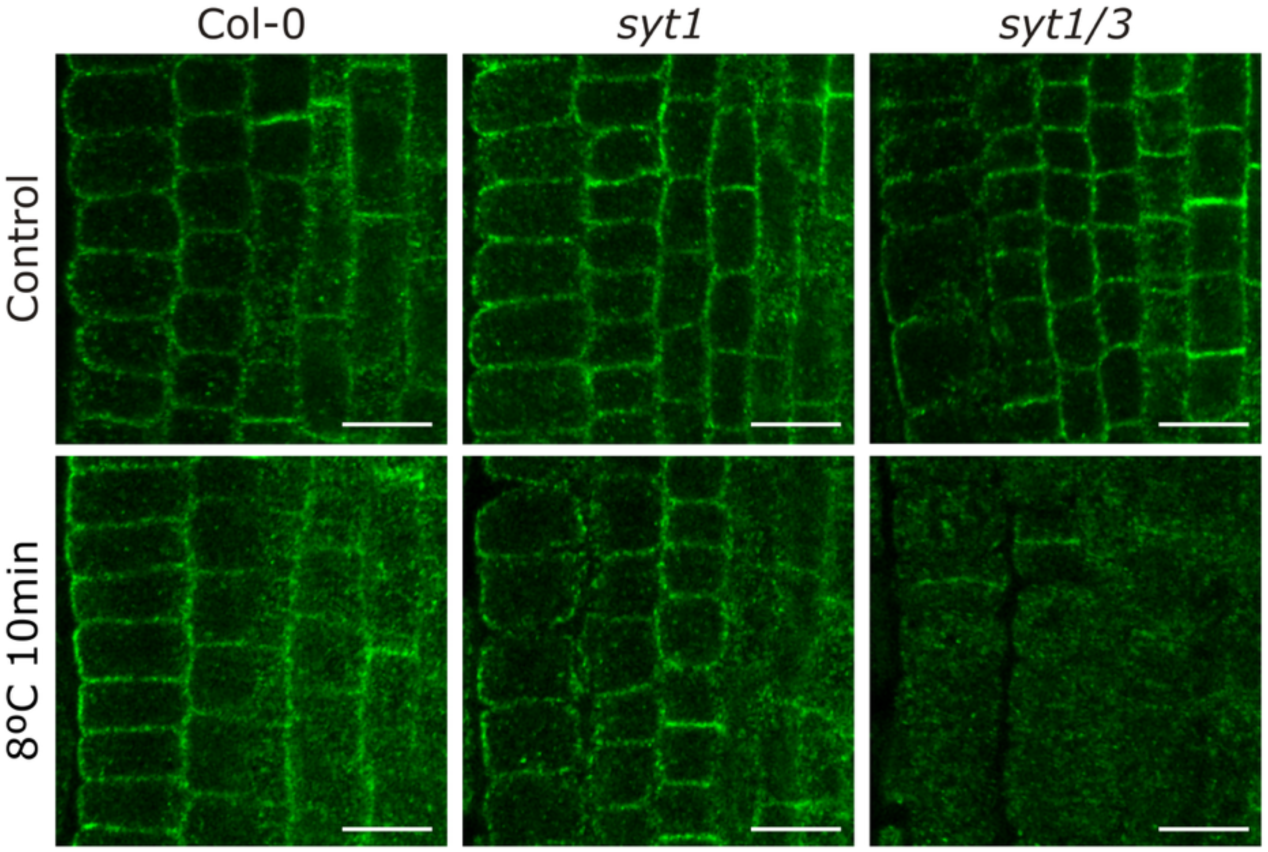
Immunolocalisation of AtEH1/Pan1 under cold stress. Representative images are shown of roots of WT (Col-0) and *syt1/3* double mutant after whole mount immunolabeling using a AtEH1/Pan1 specific antibody Plants were grown under control conditions as well as treated with cold stress. Scale bar, 10 µm.

**Figure S5.**
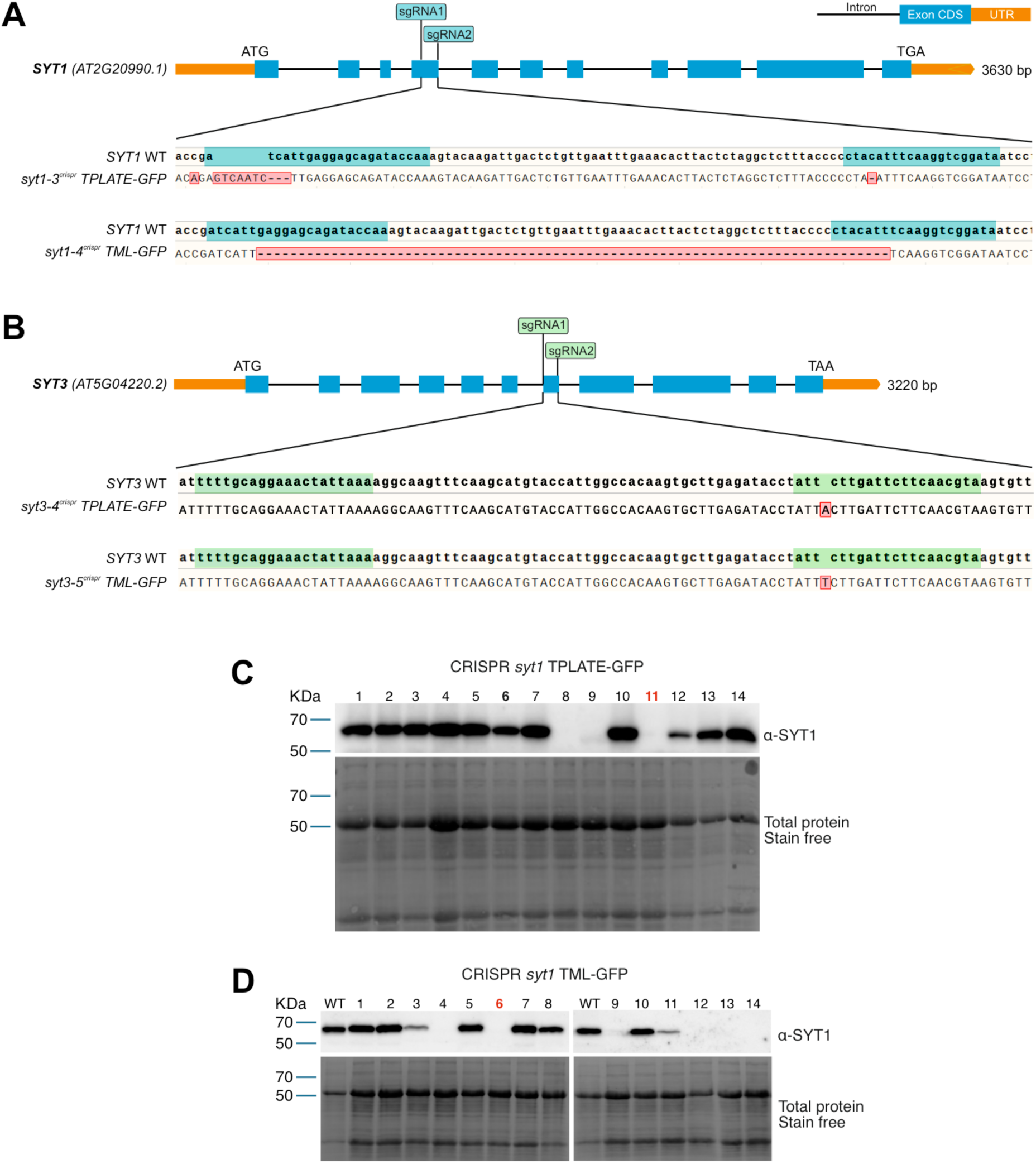
Generation and selection of CRISPR mutant lines for *SYT1* and *SYT3* in the complemented *tplate* and *tml-1* mutant backgrounds expressing TPLATE-GFP and TML-GFP rescue fusions respectively. (A and B) GuideRNAs designed for *SYT1* and SYT3 to obtain loss-of-function mutants. (C and D) Immunoblot analysis to identified *syt1* mutants generated by CRISPR in *tplate(−/−)* expressing TPLATE-GFP and *tml-1(−/−)* expressing TML-GFP background lines. The *syt1(−/−) tplate(−/−)* expressing TPLATE-GFP line 11 and *syt1(−/−) tml-1(−/−)* expressing TML-GFP line 6 were selected for further analysis are indicated in red in the immunoblot.

